# P53 Acetylation Exerts Critical Roles In Pressure Overload Induced Coronary Microvascular Dysfunction and Heart Failure

**DOI:** 10.1101/2023.02.08.527691

**Authors:** Xiaochen He, Aubrey C Cantrell, Quinesha A Williams, Wei Gu, Yingjie Chen, Jian-Xiong Chen, Heng Zeng

## Abstract

Coronary microvascular dysfunction (CMD) has been shown to contribute to cardiac hypertrophy and heart failure with preserved ejection fraction. At this point, there are no proven treatments for CMD. We have shown that histone acetylation may play a critical role in the regulation of CMD. By using a mouse model that replaces lysine with arginine at residues K98/117/161/162R of p53 (p53^4KR^), preventing acetylation at these sites, we test the hypothesis that acetylation-deficient p53^4KR^ could improve coronary microvascular dysfunction and prevent the progression of hypertensive cardiac hypertrophy and heart failure. Wild-type (WT) and p53^4KR^ mice were subjected to pressure overload (PO) by transverse aortic constriction to induce cardiac hypertrophy and heart failure (HF). Echocardiography measurements revealed improved cardiac function together with reduction of apoptosis and fibrosis in p53^4KR^ mice. Importantly, myocardial capillary density and coronary flow reserve (CFR) were significantly improved in p53^4KR^ mice. Moreover, p53^4KR^ upregulated the expression of cardiac glycolytic enzymes and glucose transporters, as well as the level of fructose-2,6-biphosphate; increased PFK-1 activity; and attenuated cardiac hypertrophy. These changes were accompanied by increased expression of HIF-1α and proangiogenic growth factors. Additionally, the levels of SERCA-2 were significantly upregulated in sham p53^4KR^ mice as well as in p53^4KR^ mice after TAC. *In vitro*, p53^4KR^ significantly improved endothelial cell (EC) glycolytic function and mitochondrial respiration, and enhanced EC proliferation and angiogenesis. Similarly, acetylation-deficient p53^4KR^ significantly improved CFR and rescued cardiac dysfunction in SIRT3 KO mice. Our data reveal the importance of p53 acetylation in coronary microvascular function, cardiac function, and remodeling, and may provide a promising approach to improve hypertension-induced coronary microvascular dysfunction (CMD) and to prevent the transition of cardiac hypertrophy to heart failure.

## Introduction

Hypertension, diabetes, and aging are the major risk factors for the development of heart failure (HF) ^1-4^, which is one of the leading causes of mortality and morbidity worldwide ^5^. Clinical studies demonstrate that coronary microvascular dysfunction (CMD), coronary capillary rarefaction, cardiac hypertrophy, and fibrosis play a detrimental role in the progression of HF ^6, 7^. Recent studies demonstrate that coronary artery remodeling and microvascular rarefaction (impairment of angiogenesis) are two important contributors to reduced coronary flow reserve (CFR) i.e. coronary microvascular dysfunction ^8-14^. CFR is an essential predictor of heart failure diseases such as diastolic dysfunction and heart failure with preserved ejection fraction (HFpEF) ^8, 9, 15, 16^. Many pathological conditions such as diabetic and hypertensive cardiac hypertrophy are associated with reduction of CFR^8, 9, 15^. Moreover, preserved CFR in diabetic patients reduces the number of cardiac events to levels seen in non-diabetic controls^17^. Accumulating evidence reveals that restoration of CFR or improvement of CMD have great promise in reducing high mortality in heart failure. At this point, there are no proven treatments for CMD due to a lack of mechanistic studies. The mechanisms related to CMD are unclear, but endothelial dysfunction and microvascular rarefaction are suggested as two important contributors to reduced CFR ^8-14^. Previous studies suggest that coronary microvascular rarefaction contributes to the progression from compensated hypertrophy to heart failure ^18-20^, whereas promotion of angiogenesis improves cardiac function and delays the progression of heart failure ^21-23^. So far, our understanding of the molecular mechanisms of microvascular rarefaction and CMD in HF is still incomplete.

Ablation of p53 has been shown to protect against coronary vascular rarefaction and cardiac injury ^24, 25^. In contrast, accumulation of p53 impaired cardiac angiogenesis and systolic function ^26^. P53-mediated impairment of cardiac angiogenesis was observed in animal models of cardiac hypertrophy^27^. Furthermore, endothelial senescence and dysfunction were associated with acetyl-p53^28^. These studies suggest that p53 plays a critical role in the development of cardiac and endothelial dysfunction. Our previous study also showed that down-regulation of Sirtuin 3 (SIRT3) was associated with increased acetylation of p53 in cardiomyocytes, whereas overexpression of SIRT3 significantly reduced p53 acetylation and blunted microvascular rarefaction, cardiac fibrosis, and hypertrophy, and improved cardiac function in diabetic mice ^29^. This study suggests that regulation of p53 activation by acetylation plays a critical role in microvascular and cardiac function. Indeed, p53 activity is finely tuned by the regulation of p53 protein stability via co-activators and inhibitors, as well as post-translational modifications, including acetylation ^30^. Acetylation of p53 plays a major role in regulating promoter-specific activation of downstream targets during stress responses ^30^. Previous study demonstrated that the acetylation-deficient p53 (K98/117/161/162) lost its ability to induce cellular senescence, cell cycle arrest, and apoptosis, and to regulate certain p53 dependent metabolic pathways in cancer cell lines^30^. However, the role of p53 acetylation in hypertensive pressure overload (PO)-induced CMD and cardiac metabolism and heart failure is largely unknown.

In this study, we examined whether acetylation-deficient p53^4KR^ preserves coronary microvascular function, prevents pathological cardiac hypertrophy and fibrosis, and the development of HF in response to chronic PO.

## Methods

The data that support the findings of this study are available from the corresponding author upon reasonable request. In addition to the section below, other detailed methods are available in the Supplemental Materials.

### Mice

Mice expressing p53 with mutations at lysine residues K98, K117, K161, and K162 (replacing lysine with arginine, p53^4KR^) mice on the C57BL/6j background were provided by Dr. Wei Gu at the Columbia University and maintained in the Laboratory Animal Facilities at the University of Mississippi Medical Center (UMMC). Male C57BL/6 mice were purchased from The Jackson Laboratory (Bar Harbor, ME) and were used as wild type (WT) controls. The SIRT3/p53^4KR^ mouse was generated by crossed SIRT3KO mouse with p53^4KR^ mouse. All animals were fed laboratory standard chow and water and housed in individually ventilated cages. All protocols were approved by the Institutional Animal Care and Use Committee of the UMMC (Protocol ID: 1564 and 1189) and were in compliance with the National Institutes of Health Guide for the Care and Use of Laboratory Animals (NIH Pub. No. 85-23, Revised 1996).

### Statistical Analysis

Data are presented as mean ± SEM. The assumption of normality in both comparison groups were determined by normality and long-normality test. Statistical significance was determined by using Student’s t-test (two-tailed) between the means of two groups, or two-way ANOVA followed by Tukey’s *post-hoc* test for multiple comparisons in GraphPad Prism 8 software (San Diego, CA). P<0.05 was considered statistically significant.

## Results

### Acetylation-deficient p53^4KR^ attenuates systolic dysfunction and hypertrophy after TAC

p53^4KR^ is a knock-in mouse model that expresses p53 with four lysine-to-arginine mutations K98/117/161/162R ^31^. These mutations abolish acetylation of p53 at these four sites, so that this form of p53 fails to regulate cellular senescence, cell cycle arrest and apoptosis, and metabolic pathways ^30^. To investigate the role of acetylation of p53 in cardiac function during PO, WT and p53^4KR^ mice were subjected to TAC for 8 weeks. Cardiac parameters measured by echocardiography are shown in supplemental Table S2. As shown in Figure 1, the expression of p53 was significantly increased in the hearts of p53^4KR^+TAC mice after TAC surgery (Figure 1A and Supplemental Figure S1A). The acetylation of p53 at K370 had a similar increase as p53 expression after TAC (Figure 1A). The ejection fraction (EF) and fractional shortening (FS) were significantly decreased in WT mice at 8 weeks after TAC when compared to the sham controls. However, the systolic function of p53^4KR^ mice subjected to TAC had no significant change (Figure 1B and Supplemental Figure S1B). The ratio of heart weight to tibia length or body weight was significantly increased in the WT mice subjected to TAC, but not in the p53^4KR^ mice (Figure 1C and Supplemental Figure S1C). TAC-induced increase of LV mass determined by echocardiography was also significantly attenuated in a p53^4KR^ mice (Figure 1D). The WT mice+TAC showed gradually decreased cardiac output over time after TAC, while p53 mutant mice+TAC had sustained cardiac output, though it was not statistically different between the two groups (Supplemental Figure S1D). The survival rate was not significantly different between WT and p53^4KR^ mice after TAC (Supplemental Figure S1E). Immunoblot further showed that the LV β-MHC protein expression was significantly attenuated in p53^4KR^ mice as compared to the WT mice subjected to TAC (Figure 1A and Supplemental Figure S1A). WGA staining also revealed that TAC caused a significantly higher increase in the cross-sectional area of cardiomyocytes in the WT mice as compared with p53^4KR^ mice (Figure 1E). Western blot analysis also revealed a significant reduction of brain natriuretic peptide (BNP) expression, a hypertrophic marker, in the heart of p53^4KR^ mice after TAC (Figure 1A). Interestingly, the basal level of BNP was also significantly reduced in the heart of p53^4KR^ mice. These data suggest that p53 acetylation deficiency ameliorates the development of cardiac hypertrophy and attenuates the deterioration of cardiac function after TAC.

**Figure 1.**
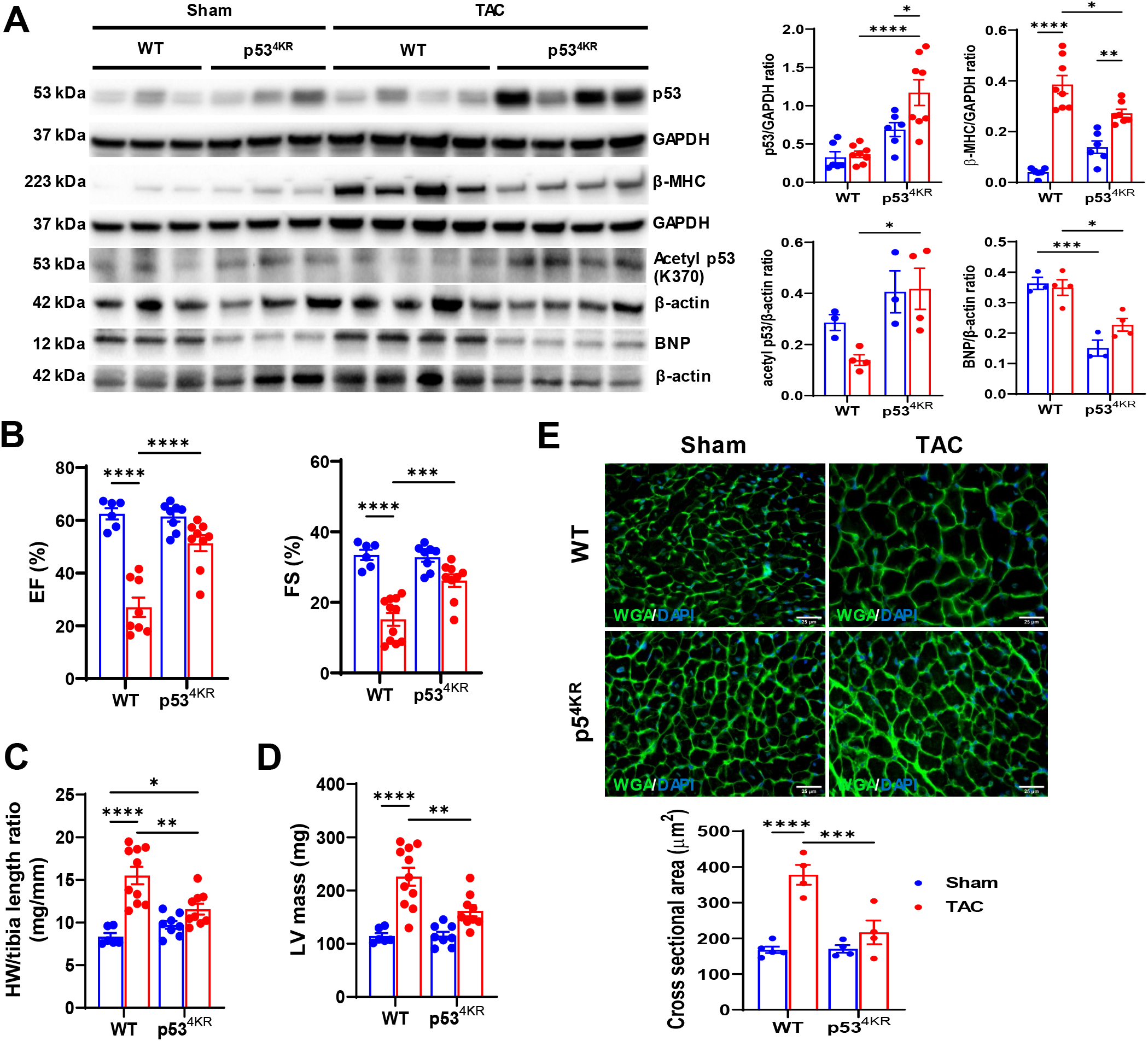
Effects of acetylation-deficient p53^4KR^ on systolic function and cardiac hypertrophy. **A**, Representative immunoblots and quantitative analysis of p53, β-myosin heavy chain (β-MHC), acetyl-p53, brain natriuretic peptide (BNP), β-actin, and GAPDH in the indicated mouse hearts subjected to either sham or transverse aortic constriction (TAC) procedure for eight weeks (n=3-4 or 6-8). **B**, Left ventricular (LV) ejection fraction (EF) and fractional shortening (FS) measured by echocardiography in the indicated groups (n=6-10). **C**, Ratio of heart weight to tibia length in the indicated groups (n=6-10). **D**, LV mass measured and calculated by echocardiography in the indicated groups (n=6-10). **E**, Representative images of wheat germ agglutinin–stained frozen heart sections in the indicated groups. (n=4–5). A minimum of 100 cardiomyocytes from each LV section of each mouse was measured. Bar=25 μm. \**p*<0.05, \*\**p*<0.01, \*\*\**p*<0.001, \*\*\*\**p*<0.0001.

### Acetylation-deficient p53^4KR^ preserves diastolic function and reduces cardiac apoptosis and fibrosis after TAC

We further investigated the role of p53 acetylation deficiency on the diastolic function during PO-induced heart failure. Doppler measurements are shown in supplemental Table S2. WT mice subjected to TAC exhibited a dramatic increase in E/A ratio (greater than 2, 2.05±0.10 versus 1.32±0.07 in the sham control) and E/e’ ratio (Figure 2A and Supplemental Figure S2A), suggesting elevation of left atrial pressure and development of diastolic dysfunction when compared to the p53^4KR^ mice after TAC. Western blot and immunostaining analysis showed that the expression of SERCA-2 ATPase was significantly upregulated in p53^4KR^ mice at basal levels and after TAC (Figure 2C and D). Additionally, interstitial fibrosis was significantly increased in WT mice (but not in p53^4KR^ mice) after TAC (Figure 2B). Platelet-derived growth factor β-receptor (PDGFR-β) and fibroblast specific protein 1 (FSP1) regulate tissue fibroblast activation. Interestingly, cardiac PDGFR-β and FSP-1 protein expression were significantly attenuated in p53^4KR^ mice as compared with WT mice after TAC (Figure 2C and Supplemental Figure S2B). p53^4KR^ mice had low basal levels of collagen-1 in the heart. Similarly, cardiac collagen-1 levels were significantly reduced in p53^4KR^ mice as compared with WT mice after TAC (Figure 2C and Supplemental Figure S2B), suggesting that reduced cardiac fibroblast activation may be involved in the attenuated fibrosis observed in p53^4KR^ mouse hearts. In addition, fewer TUNEL positive cells were detected in hearts of p53^4KR^ mice compared with WT mice after TAC (Figure 2E), indicating that p53 acetylation deficiency attenuated cardiac apoptosis in mice after TAC. To understand which cells are undergoing apoptosis, cells were co-stained for apoptotic marker TUNEL along with either cardiomyocyte marker Troponin-T, or myofibroblast marker FSP-1. Our data showed that the TUNEL positive cells predominantly cardiomyocytes (Supplemental Figure S2C). In addition, there were a few TUNEL positive myofibroblast cells after TAC (Supplemental Figure S2D).

**Figure 2.**
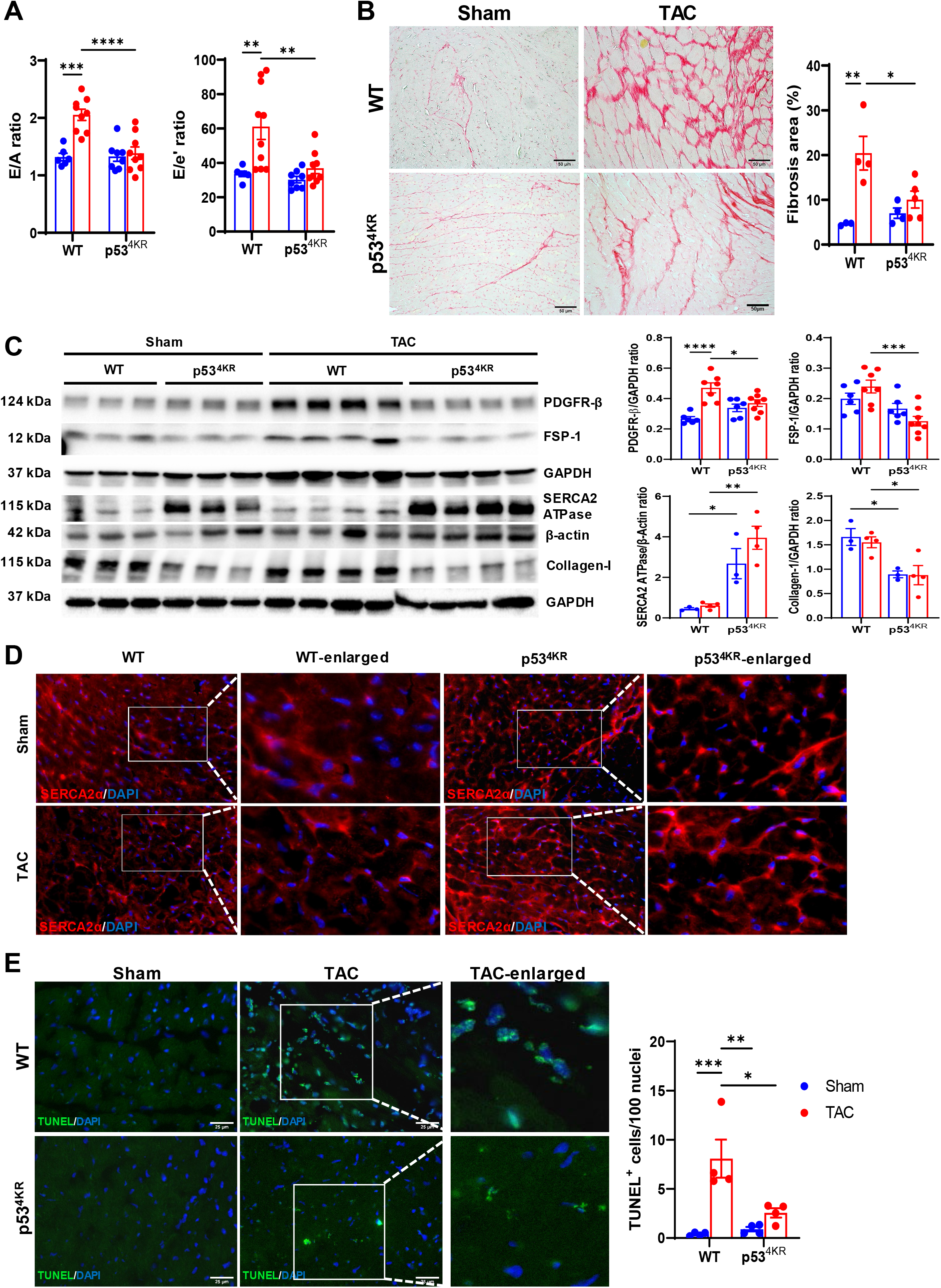
Effects of acetylation-deficient p53^4KR^ on diastolic function, cardiac fibrosis, and apoptosis. **A**, Representative pulsed-wave Doppler and tissue Doppler images from an apical 4-chamber view of WT and p53^4KR^ mice subjected to either sham or TAC procedure for eight weeks and ratio of the peak velocity of early (E) to late (A) filling of mitral inflow (E/A) in the indicated groups (n=6-9). The ratio of E to the tissue motion velocity in early diastole (e’) was calculated in the indicated groups (n=6-10). **B**, Representative images of Picrosirius red-stained paraffin-embedded heart sections and quantification of the percentage of interstitial fibrosis area in the indicated groups (n=3-5). Bar=50 μm. **C**, Representative immunoblots and quantitative analysis of PDGFR-β, FSP-1, collagen-1, SERCA2 ATPase, β-actin, and GAPDH in the indicated mouse hearts (n=3-4 or 6-8). **D**, Representative images of staining of SERCA2α and DAPI in frozen heart sections. **E**, Representative images of TUNEL-stained frozen heart sections and quantification of the number of TUNEL-positive cells (green)/field in the indicated groups (n=4). 5-15 randomly selected fields per LV section of each mouse was measured. Bar=25 μm. \**p*<0.05, \*\**p*<0.01, \*\*\**p*<0.001, \*\*\*\**p*<0.0001.

### Acetylation-deficient p53^4KR^ preserves capillary density and coronary flow reserve in heart failure

Deletion of p53 has been shown to promote angiogenesis ^24^. To determine whether p53 acetylation deficiency is able to stabilize the cardiac microvasculature during PO, isolectin B4-positive area (Figures 3A) was quantified. The result shows a significant decrease of cardiac capillary density in WT mice after TAC, while cardiac capillary density was preserved in p53^4KR^ mice after TAC. p53-mediated hypoxia-independent HIF-1α degradation is involved in angiogenesis inhibition following PO ^26^. We further explored the molecular mechanism by which p53^4KR^ regulates angiogenic signaling pathways. Cardiac HIF-1α protein expression was significantly elevated in p53^4KR^ mice but not in WT mice after TAC (Figure 3B and Supplemental Figure S3A). To identify where HIF-1α activation was taking place, the expression of HIF-1α in the cardiomyocytes, endothelial cells and fibroblasts were examined by co-staining HIF-1α with cardiomyocyte marker Troponin-T, endothelial marker IB4, or myofibroblast marker FSP-1. Immunohistochemical study revealed that expression of HIF-1α was primarily localized to the cardiomyocyte, whereas endothelial cells and myofibroblasts also showed a few positive cells (Figure 3C and Supplemental Figure S3B and S3C). Cardiac angiopoietin-1 protein expression was increased in p53^4KR^ mice as compared with WT mice under control conditions, and the expression was further increased in p53^4KR^ after TAC (Figure 3B). TAC did not affect cardiac angiopoietin-1 protein expression in WT mice (Figure 3B and Supplemental Figure S3A). In addition, cardiac VEGF protein expression was markedly reduced in WT mice after TAC, while cardiac VEGF protein expression was unchanged in p53^4KR^ mice (Figure 3B and Supplemental Figure S3A). Moreover, the expression of heme oxygenase-1 (HO-1), a direct p53 target with anti-inflammatory, antioxidant, antiapoptotic, and proangiogenic effects, was significantly elevated in p53^4KR^ mice after TAC (Figure 3B and Supplemental Figure S3A). Cardiac VCAM-1 expression was significantly increased in WT but not in p53^4KR^ mice after TAC (Figure 3B and Supplemental Figure S3A), suggesting that endothelial inflammatory activation was attenuated in p53^4KR^ mice after TAC. These data suggest that p53 acetylation deficiency preserved cardiac angiogenesis, and attenuated endothelial activation after TAC.

**Figure 3.**
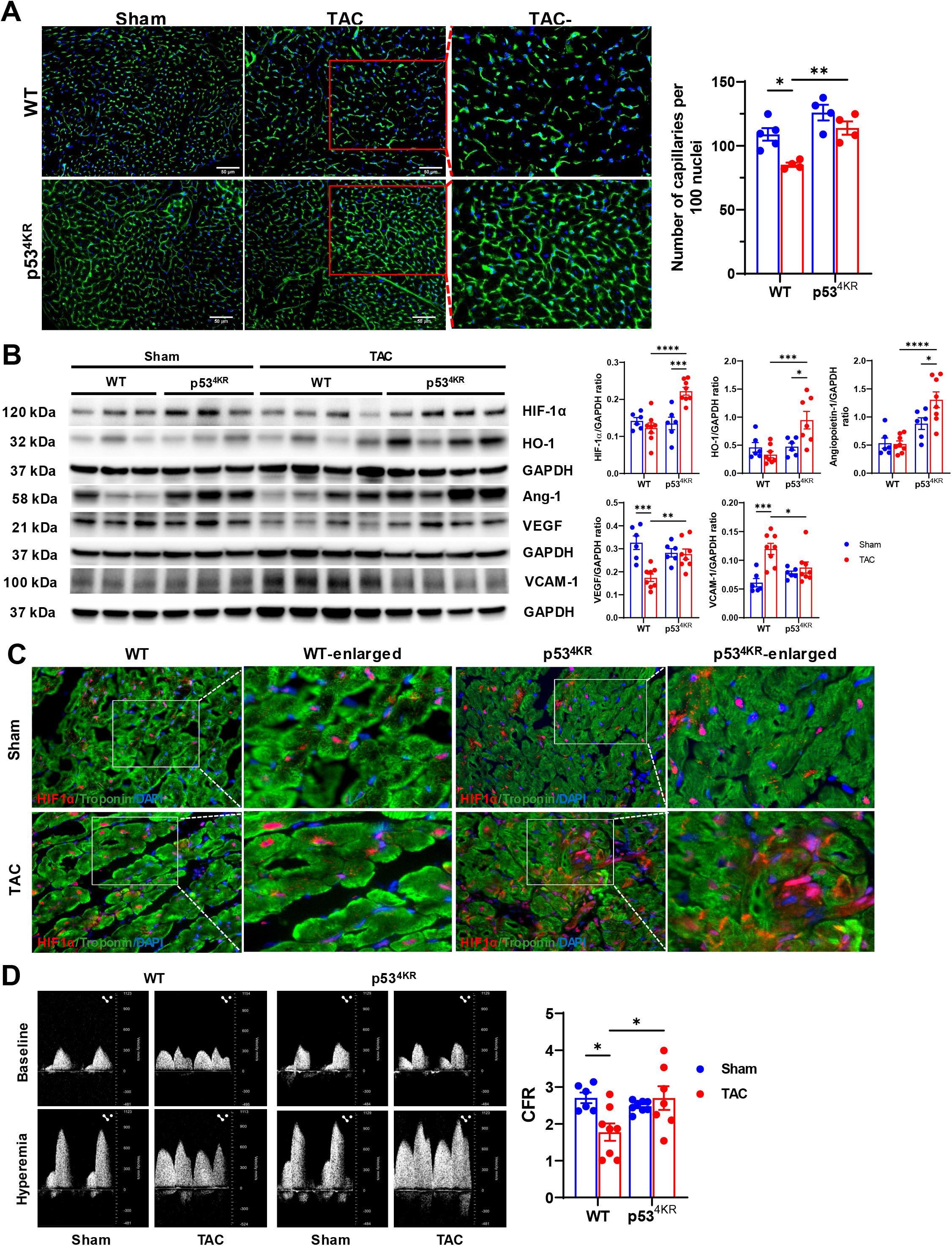
Acetylation-deficient p53^4KR^ preserves coronary vasculature and coronary flow reserve (CFR). **A**, Representative images of Isolectin B4 (IB4, green; DAPI stains the nuclei, blue)-stained frozen heart sections and quantification of the number of capillaries/100 nuclei in the indicated groups (n=4-5). Bar=50 μm. **B**, Representative immunoblots and quantitative analysis of HIF-1α, HO-1, Ang-1, VEGF, VCAM-1, and GAPDH in the indicated mouse hearts (n=6-8). **C**, Representative images of co-staining of cardiomyocyte marker Troponin-T, endothelial marker IB4, and myofibroblast marker FSP-1 with HIF-1α in frozen heart sections. **D**, Representative pulsed-wave Doppler images of the proximal left coronary arteries and CFR of WT and p53^4KR^ mice subjected to either sham or TAC procedure for eight weeks. CFR was calculated as the ratio of hyperemic peak diastolic flow velocity (2.5% isoflurane) to baseline peak diastolic flow velocity (1% isoflurane) in the indicated groups (n=6-8). \**p*<0.05, \*\**p*<0.01, \*\*\**p*<0.001, \*\*\*\**p*<0.0001.

Loss of capillary or microvascular rarefaction is one of the key contributors to the development of coronary microvascular dysfunction; therefore, we further examined whether loss of acetylation of p53 protects microvascular function in PO-induced heart failure. We measured the coronary flow velocity (Supplemental Table S2) at 1% (baseline) or 2.5% isoflurane (hyperemia) to calculate the CFR. As shown in Figure 3D, PO significantly reduced the CFR in the WT mice, while CFR was preserved in the p53^4KR^ mice.

### P53 acetylation deficiency elevates glycolysis related enzyme and glucose transporters

To explore the molecular events by which p53^4KR^ regulates cardiac remodeling, we determined the expression levels of some glycolysis-related proteins. Cardiac PFK-1 expression was significantly elevated in p53^4KR^ mice after TAC, while its expression was unchanged in WT mice (Figure 4A and Supplemental Figure S4A). In addition, the expression of glucose transporter Glut-1 was reduced similarly in both WT and p53^4KR^ mice after TAC, but it was significantly higher in the p53^4KR^ mice after TAC (Figure 4A and Supplemental Figure S4A). Cardiac Glut-4 expression was significantly elevated in p53^4KR^ mice after TAC, but its expression was not affected in WT mice after TAC (Figure 4A and Supplemental Figure S4A).

**Figure 4.**
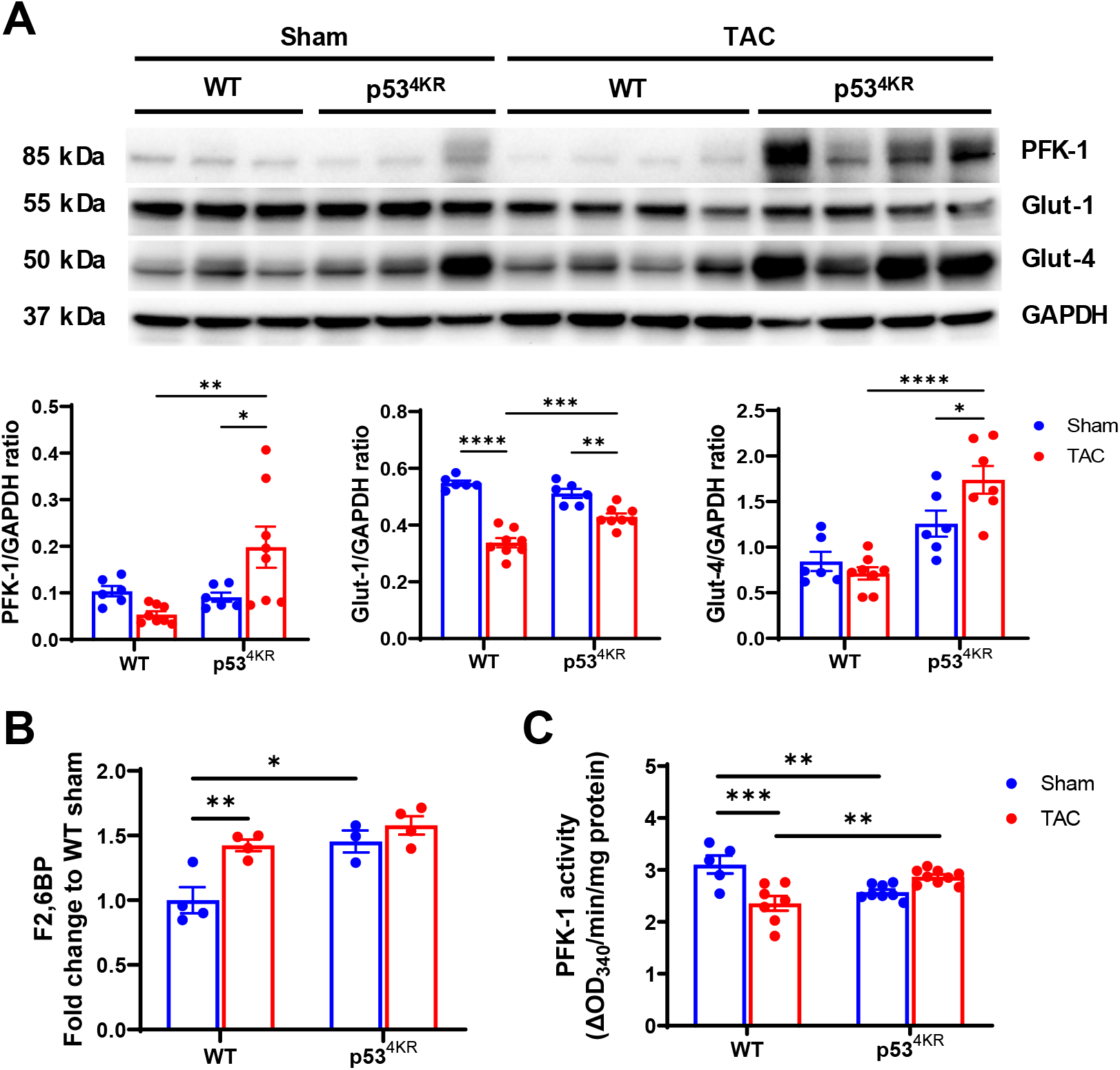
Acetylation-deficient p53^4KR^ increases cardiac glucose transporters and glycolytic function. **A**, Representative immunoblots and quantitative analysis of PFK-1, GLUT-1, GLUT-4, and GAPDH in the indicated groups. **B**, Cardiac F2,6-BP level was determined by the coupled-enzymatic assay and expressed as the fold change to the WT sham group. n=3-4. **C**, Cardiac PFK-1 activity was determined by the coupled-enzymatic assay and expressed as the OD_340nm_/min/mg protein. n=5-9. \**p*<0.05, \*\**p*<0.01, \*\*\**p*<0.001, \*\*\*\**p*<0.0001.

By using coupled-enzymes methods, we measured the level of glycolytic intermediate F2,6-BP, as well as PFK-1 activity. The basal level of F2,6-BP in the hearts of p53^4KR^ mice was significantly higher than the WT mice (Figure 4B). WT mice subjected to TAC exhibited a significant elevation in the level of F2,6-BP, whereas no change was observed in p53^4KR^ mice after TAC (Figure 4B). Moreover, PFK-1 activity in the WT mice after TAC was significantly reduced when compared to the sham controls, whereas it was significantly increased in p53^4KR^ mice after TAC (Figure 4C). These data suggest that p53 acetylation deficiency enhanced glycolytic function by increasing the level of F2,6-BP and enhancing PFK-1 activity.

### P53 acetylation deficiency increases glycolysis in the endothelial cell (EC) and aortic sprouting

Since p53^4KR^ improved glycolytic function and increased expression of cardiac proangiogenic growth factors, we further investigated whether p53^4KR^ enhanced glycolysis and angiogenesis in isolated ECs from WT and p53^4KR^ aortas. Cell proliferation was significantly increased in mouse aortic endothelial cells (MAECs) from p53^4KR^ mice as compared with MAECs from WT mice (Figure 5A). Moreover, p53^4KR^ ECs exhibited a significant upregulation of basal glycolysis, glycolytic capacity, and glycolytic reserve when compared to WT ECs (Figure 5B). This was associated with a significant increase in mitochondrial basal respiration and ATP production (Figure 5C). In addition, the vessel explant sprouting study demonstrated a significant increase in the vessel explant sprouting area in the vessel explants of p53^4KR^ mice as compared to the WT mice (Figure 5D). These data suggest that p53 acetylation deficiency can also promote endothelial glycolysis and *ex vivo* angiogenesis.

**Figure 5.**
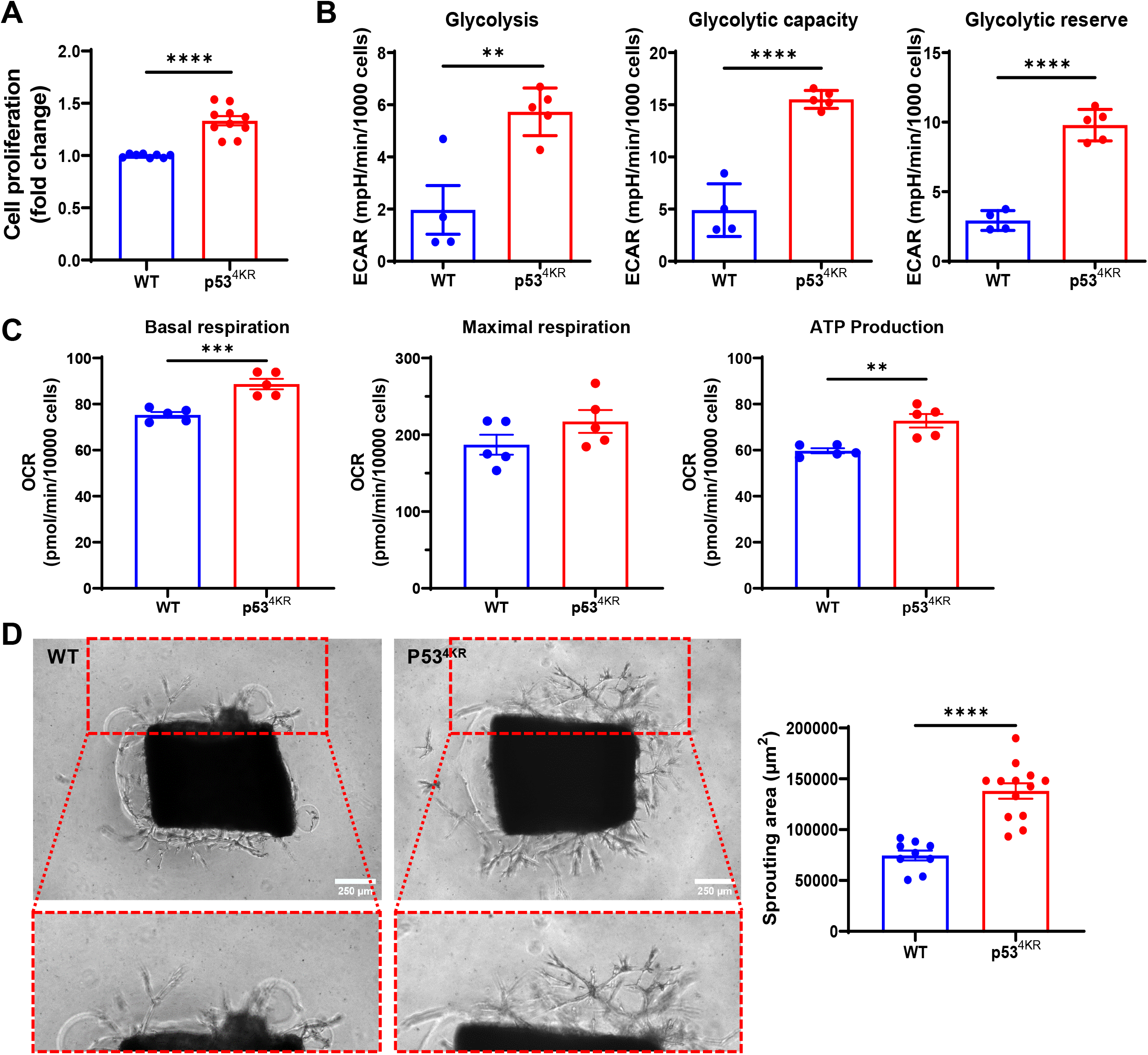
Acetylation-deficient p53^4KR^ increases glycolysis in MAECs and aortic sprouting. **A**, Cell proliferation of p53^4KR^ MAECs was significantly higher than that of the WT MAECs, as measured by MTT assay. n=8-10. **B**, Glycolysis stress test and extracellular flux analysis of glycolysis, glycolytic capacity, and glycolytic reserve in MAECs isolated from p53^4KR^ and WT mice. n=4-5. **C**, Mitochondrial stress test and extracellular flux analysis of mitochondrial basal and maximal respiration and ATP production in MAECs isolated from p53^4KR^ and WT mice. n=5. **D**, Representative images of the aortic ring sprouting assay at day 5 of incubation and quantification of the sprouting area in the indicated groups (n=9-13). Bar=250 μm. \*\**p*<0.01, \*\*\**p*<0.001, \*\*\*\**p*<0.0001.

### Acetylation-deficient p53^4KR^ preserves coronary flow reserve and rescues heart failure in SIRT3KO mice

To validate the p53^4KR^-dependent rescue of hypertension-induced CMD, cardiac hypertrophy and heart failure, the genetic SIRT3 knockout (SIRT3 KO) mouse was crossed with p53^4KR^ to generate a SIRT3KO/p53^4KR^ mouse. As shown in Figure 6, knockout of SIRT3 resulted in the impairment of CFR (Figure 6A), along with heart failure as evidenced by reductions in EF% and FS% (Figure 6B and Supplemental Figure S4B), and elevations in IVRT, MPI and E/e’ ratio (Figure 6C). Consistent with our TAC heart failure model, p53 acetylation deficiency significantly improved CFR and rescued cardiac diastolic dysfunction and heart failure in SIRT3KO mice.

**Figure 6.**
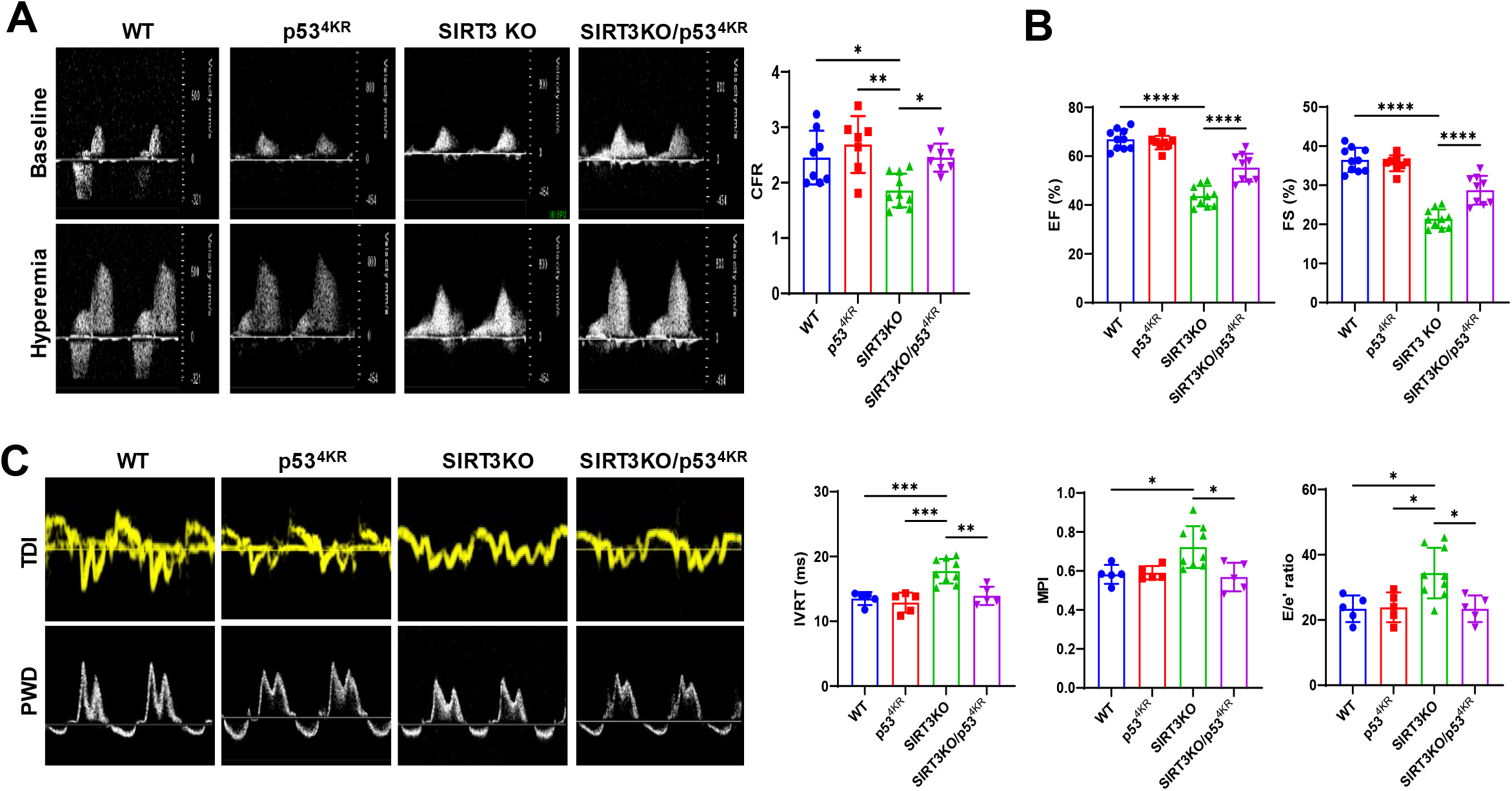
Effects of acetylation-deficient p53^4KR^ on CFR and cardiac dysfunction in SIRT3KO mice. **A**, Representative pulsed-wave Doppler images of the proximal left coronary arteries and CFR of WT, SIRT3KO, p53^4KR^ and p53^4KR^/SIRT3KO mice. CFR was calculated as the ratio of hyperemic peak diastolic flow velocity (2.5% isoflurane) to baseline peak diastolic flow velocity (1% isoflurane) in the indicated groups (n=7-11). **B**, Left ventricular (LV) ejection fraction (EF) and fractional shortening (FS) measured by echocardiography in the indicated groups (n=8-10). **C**, Representative pulsed-wave Doppler and tissue Doppler images from an apical 4-chamber view in the indicated groups. The diastolic function isovolumic relaxation time (IVRT), myocardial performance index (MPI), and ratio of E to the tissue motion velocity in early diastole (e’) was calculated in the indicated groups (n= 5-9). \**p*<0.05, \*\**p*<0.01, \*\*\**p*<0.001, \*\*\*\**p*<0.0001.

## Discussion

In this study, we examined the effect of acetylation-deficient p53 on coronary microvascular dysfunction, cardiac hypertrophy, and the development of HF in response to chronic PO. The main findings are that p53 acetylation deficiency prevents coronary capillary rarefaction and improves CFR while attenuating cardiac remodeling and dysfunction. These changes were associated with enhanced cardiac glycolytic function and increased levels of proangiogenic growth factors, involving PFK-1, glucose transporters, HIF-1α, Ang-1, VEGF, and HO-1. Thus, our data support the importance of p53 acetylation in coronary microvascular dysfunction and cardiac function, as well as cardiac remodeling during PO-induced HF.

The tumor suppressor protein p53 plays a critical role in the pathogenesis of heart disease, as evidenced by the increased level of p53 and the number of apoptotic cells in patients with heart disease ^32^. p53 regulates the cellular stress response to DNA damage, oxidative stress, hypoxia, or cytokines, which induces expression of p53 target genes that modulate cell cycle and growth arrest, as well as apoptosis^33^. Thus, by using experimental animals with global, cardiomyocyte-specific, or endothelial-specific p53 gene deletion, as well as pharmacological inhibition of p53, studies have demonstrated that inhibition of p53 effectively attenuated TAC or myocardial infarction (MI)-induced myocardial apoptosis, ventricular remodeling, vascular rarefaction, and heart failure ^24-26^. In addition to the classical regulation of p53 stability by Mdm2, accumulating studies suggest that p53 activity is also rapidly and effectively regulated by post-translational modification, such as phosphorylation and acetylation. Specifically, acetylation of p53 plays a major role in controlling its binding to the promoter region of specific downstream targets during stress responses. Our recent study demonstrated that reduced p53 acetylation was associated with improved microvascular angiogenesis and cardiac function in diabetic mice, exhibiting decreased cardiac fibrosis and hypertrophy ^29^. A study by Wang *et al* demonstrated that cells expressing the acetylation-deficient p53^4KR^ mutant were protected from apoptosis, growth arrest, and ferroptosis^30^. The same effect was observed in p53^4KR^ mice that were tumor prone but failed to develop early-onset tumors ^31^. Our present study demonstrated a significant reduction of apoptosis, although the p53 level was increased in the heart of p53^4KR^ mice subjected to PO. This result indicates a crucial role of p53-acetylation in mediating apoptosis independent of p53 level, which is consistent with the previous study ^30^. Our results show that acetylation of p53 at K370 is upregulated in p53^4KR^ + TAC mice relative to WT+TAC and p53^4KR^ sham mice, while WT+TAC mice are unchanged relative to WT+ sham. We suspect that the acetylation of p53 at K370 may be upregulated in the p53^4KR^+TAC mice as a compensatory mechanism, whereby the cell is trying to increase acetylated p53 at one or more of these sites by increasing translation of p53 itself in order to respond to the TAC and mitigate the damage caused. This would suggest that the acetylation deficiency at these residues is causing reduction, if not complete loss, of function for acetylated p53 in this capacity. Additionally, as previously demonstrated by Wang *et al*., loss of acetylation at K117 in mice reduced the ability of p53 to mediate apoptosis, while it was still capable of inducing cell senescence and cell cycle arrest ^30^. By mutating two additional lysine residues (K161 and K162) they found that p53 could no longer regulate cell senescence or cell cycle arrest. Furthermore, Kon *et al*. found that adding a mutation at the K98 residue abolishes p53-mediated ferroptosis, which is retained in models without this mutation ^31^. They further identified that the mTOR pathway is intact in p53^4KR^ mice and generated an additional mutation at K136 to create a p53^5KR^ model which has impaired mTOR pathway activity. Therefore, we believe that individual acetylation sites may regulate different physiological activities of p53, and that altering which acetylation sites are mutated will impact the functional abilities of the p53 protein. While our p53^4KR^ model has diminished function, it does not inhibit every pathway by which p53 acts, although these 4 sites are critical mediators of various pathways and as such are important for p53 activity. Previous study demonstrated a critical role of p53 acetylation in ferroptosis in which p53^4KR^ further lost its function to induce ferroptosis when compared to the WT-p53 or p53^3KR^ (K117/161/162R) in cancer cell lines ^30^. We have not found that PO-induced HF was correlated with myocardial ferroptosis at 8 weeks after TAC. Although HO-1 was upregulated, the level of glutathione peroxidase 4 and 4-hydroxynonenal were not altered in mouse hearts after PO at this time point (He et al unpublished data). These data suggest that acetylation-deficient p53^4KR^ functions in mediating apoptosis and ferroptosis in the heart, and thus protects cardiac function.

One of the key feature of pressure overload-induced heart failure is microvascular rarefaction and impairment of CFR. A recent study also revealed that pressure overload induces major transcriptional and metabolic adaptations in cardiac microvascular endothelial cells (MiVEC) resulting in excess interstitial fibrosis and impaired angiogenesis. Our present study showed a very similar cardiac microvascular rarefaction, with reduced coronary blood flow in pressure overload-induced heart failure. These studies highlight the critical role of microvascular rarefaction and MiVEC as key pathogenic modulators of pressure overload–induced heart disease in mice, suggesting new therapeutic targets to alter angiogenesis in pressure overload– induced cardiac remodeling. Global p53 deficiency or systemic p53 inhibition by pifithrin-α has been shown to increase cardiac angiogenesis and protect the heart from PO- or MI-induced heart failure ^25, 26^. In addition, endothelial p53 deletion improved myocardial angiogenesis and attenuated cardiac fibrosis, remodeling, and dysfunction ^24^. In line with these previous findings, our study revealed that p53^4KR^ acetylation deficiency preserved myocardial capillary density and cardiac contractile function, and attenuated cardiac hypertrophy and fibrosis. Endothelial glycolytic function significantly contributes to angiogenic growth factor, VEGF, induced angiogenesis. ECs generate most ATP through glycolysis to drive angiogenesis ^34^. Extracellular flux analysis demonstrates that glycolysis, glucose capacity, and glycolytic reserve were significantly increased in the p53^4KR^ MAECs and that aortic spouting was enhanced in the p53^4KR^ aortas compared to the WT mice. Most importantly, p53^4KR^ improved coronary vascular function as evidenced by increased CFR and preserved diastolic function. CFR is an independent determinant of long-term cardiovascular events, acute coronary syndrome, HF, and cardiac mortality in patients with coronary artery disease ^35, 36^. Impaired CFR is associated with increased myocardial infarction size, reduced LV ejection fraction, adverse LV remodeling, and reduced long-term survival in animal studies ^37^. Our previous studies have demonstrated the impaired CFR is caused, at least in part, by coronary vascular rarefaction due to reduced endothelial glycolytic function, proangiogenic, and HIF signaling in SIRT3KO mice ^37-41^. Furthermore, knockout of SIRT3 leads to a significant increase in acetylation of p53 in mouse hearts. Most importantly, cardiomyocyte-specific knockout of SIRT3 causes mitochondrial-specific lysine acetylation and p53 acetylation, which does not occur in the cytosol (Cantrell AC, unpublished study). Our present study showed that acetylation-deficient p53 significantly improves CFR and rescues the heart failure phenotype of SIRT3 KO mice. These data suggest a critical role of p53 acetylation in the mitochondria, where SIRT3 is present. Impaired CFR will lead to insufficient perfusion to meet the metabolic demand, causing hypoxia in the heart and promoting the transition from adaptive to maladaptive cardiac remodeling ^42^. Cardiac p53 has been shown to mediate HIF-1α degradation independent of hypoxia following PO ^26^. Our data showed that HIF-1α and Ang-1 levels were significantly elevated in p53^4KR^ mice, whereas no changes were observed in WT animals. Also, the level of VEGF was significantly reduced in WT mice, but was preserved in p53^4KR^ mice. Interestingly, HO-1 was identified as a direct p53 target gene that possesses potent proangiogenic, anti-inflammatory, antioxidant, and antiapoptotic effects ^43, 44^. p53^4KR^ mice exhibited significantly increased expression of HO-1, along with increased p53 levels, suggesting p53 acetylation deficiency can still regulate the expression of HO-1. Moreover, the level of VCAM-1 was upregulated in WT mice after TAC, whereas no change was observed in p53^4KR^ mice, suggesting that PO caused an increased inflammatory response in the vascular endothelium which can be suppressed by p53^4KR^.

Glucose metabolism plays an important role in diastolic function in heart disease ^45^. A metabolic shift from fatty acid oxidation to glycolysis in cardiomyocytes has been reported during PO-induced cardiac remodeling and diastolic dysfunction ^46, 47^. In addition, inhibition of glycolysis caused greater impairment of diastolic function ^48, 49^, whereas increased glycolytic substrates protects against diastolic dysfunction ^50^. Previous studies demonstrated that decreased F2,6-BP level and PFK-1 activity resulted in more profound cardiac hypertrophy, fibrosis, and dysfunction in response to PO ^51, 52^. Glucose uptake via glucose transporters also plays a critical role in regulating glycolysis, cardiac hypertrophy, and diastolic function ^53, 54^. Wild type p53 has been shown to possess a repressive effect on transcriptional activity of the GLUT1 and GLUT4 gene promoters ^55^. In the present study, p53^4KR^ seemed to lose its function to suppress GLUT-1, GLUT-4, and possibly PFK-1, which led to the increased or preserved expression of these proteins. Moreover, coupled-enzyme methods showed that the level of F2,6-BP and PFK-1 activity were significantly increased. Cardiac fibrosis is one of the key factors that contribute to the development of diastolic dysfunction ^3, 56-58^. Histological analysis demonstrated that fibrosis was significantly reduced in p53^4KR^ mice along with the levels of PDGFR-β and FSP-1, suggesting that p53^4KR^ mutations may also preserve the ability of p53 to repress PDGFR-β signaling ^59^ and fibroblast activation. Taken together, these data suggest p53^4KR^ attenuated PO-induced diastolic dysfunction, at least in part by enhancing cardiac glucose utilization via inducing PFK-1 and glucose transporters, and attenuated cardiac fibrosis.

Calcium ion (Ca^2+^) cycle plays a critical role in the contraction and relaxation of cardiomyocytes. Sarcoplasmic/endoplasmic reticulum Ca^2+^ ATPase 2 (SERCA2 ATPase) functions as a calcium transporter responsible for reuptake of Ca^2+^ by the sarcoplasmic reticulum (SR) during the contraction cycle of the heart, resulting in the myocardial relaxation that characterizes diastole. In heart failure, SERCA2 ATPase has been found to have impaired activity, causing diastolic dysfunction and diminished myocardial relaxation, while increasing levels of SERCA2 ATPase has been shown to improve heart function to some degree ^60^. During pressure overload, cardiac-specific overexpression of SERCA2a attenuated cardiac hypertrophy and improved contractility in rodents ^61, 62^, while inhibition of SERCA2 ATPase by sarcolipin impaired cardiac function in mice treated with isoproterenol. Here, we found for the first time that SERCA2 ATPase is significantly increased in p53^4KR^ mice compared with WT mice, as well as in p53^4KR^ + TAC mice compared with WT + TAC. This is likely a contributing factor to the improved cardiac function that p53^4KR^ mice demonstrate under TAC conditions, as the increased levels of SERCA2 ATPase would cause more calcium reuptake by the SR and thus improve cardiac relaxation during diastole. Although wild-type p53 has been shown to bind to and activate SERCA2 ATPase at the ER^63^, for the first time we showed that p53 is involved in regulating the protein level of SERCA2 ATPase in the heart, and that p53^4KR^ still retained that ability.

In conclusion, our findings suggest that acetylation-deficient p53 failed to suppress HIF-1α and PDGFR-β signaling, PFK-1 activity, and transcription of glucose transporters during PO-induced HF, but it can still induce SERCA2 ATPase, HO-1, proangiogenic growth factors, and endothelial anti-inflammatory signaling, despite the loss of its pro-apoptotic function. All of these contribute to myocardial angiogenesis, improved CFR, and attenuated cardiac dysfunction and remodeling. Thus, our data support the importance of p53 acetylation in CMD and cardiac function and may provide a promising approach to improve coronary vascular function and PO-induced cardiac remodeling in order to prevent the transition to heart failure.

## Funding

This work was supported by the National Heart, Lung, and Blood Institute (R01HL102042, JX Chen) and National Institute of General Medical Sciences and National Heart, Lung, and Blood Institute (R01HL151536, JX Chen), the National Institute of General Medical Sciences of the National Institutes of Health under Award Number P20GM104357 (H.Z), the National Heart, Lung, and Blood Institute (R56HL164321, H.Z).

## Contributions

X. He, H. Zeng, and JX. Chen designed the research; X. He and H. Zeng, AC. Cantrell, QA Williams performed the research and analyzed the data; W. Gu developed the mouse model and revised manuscript; X. He and JX. Chen wrote the paper. X. He, AC. Cantrell, QA Williams, W. Gu, YJ. Chen, H. Zeng, and JX. Chen revised manuscript. All authors read and approved the final manuscript

## Ethics declarations

### Ethics approval

No human tissues or samples were used. All protocols were approved by the Institutional Animal Care and Use Committee of the UMMC (Protocol ID: 1564 and 1189) and were in compliance with the National Institutes of Health Guide for the Care and Use of Laboratory Animals (NIH Pub. No. 85-23, Revised 1996).

### Conflict of interest

The authors declare no competing interests.

## Notes

### Competing Interest Statement

The authors have declared no competing interest.

